# Quantifying Chemical Fluxes Underlying Gut Microbiota–Host Interactions

**DOI:** 10.64898/2026.09.27.754802

**Authors:** Jonas Cremer, Markus Arnoldini

**Affiliations:** Department of Biology, Stanford University, Stanford, CA, USA; Department of Health Science and Technology, ETH Zürich, Zürich, Switzerland

## Abstract

To resolve how the gut microbiota shapes human health, it is essential to quantify the exchange of chemicals between microbes and the host. Much as the dose of a drug determines its pharmacological effect, the magnitude of a specific chemical flux determines its impact. Here, we establish from first principles how fluxes can be estimated by integrating key parameters from human digestive physiology and microbial metabolism. We apply this framework to three cases: the exchange of fermentation products, the production of toxins by bacterial pathogens, and nitrogen homeostasis. Our analysis identifies three directly measurable host-level parameters which, in concert with microbial activity, shape flux magnitudes: intestinal transit time, absolute microbial abundance in feces, and fecal mass loss. The simultaneous quantification of these parameters provides a feasible yet essential step toward a mechanistic and quantitative description of gut microbiota–host interactions across health and disease.

## 1. Introduction

The diverse microbial community of the human gut has profound impacts on host physiology and health [1, 2]. Microbiota–host interactions are largely mediated by the exchange of a broad range of chemicals, including short-chain fatty acids [3], neurotransmitters [4, 5], vitamins [6], and toxins [7] (Figure 1A). Potential positive and negative health effects have been identified for many of these molecules. However, to determine if and how a specific molecule matters in a given health context, it is essential to quantify how much of it is exchanged per unit time. In other words, we need to quantify the fluxes of chemicals between the microbiota and the host.

**Figure 1:**
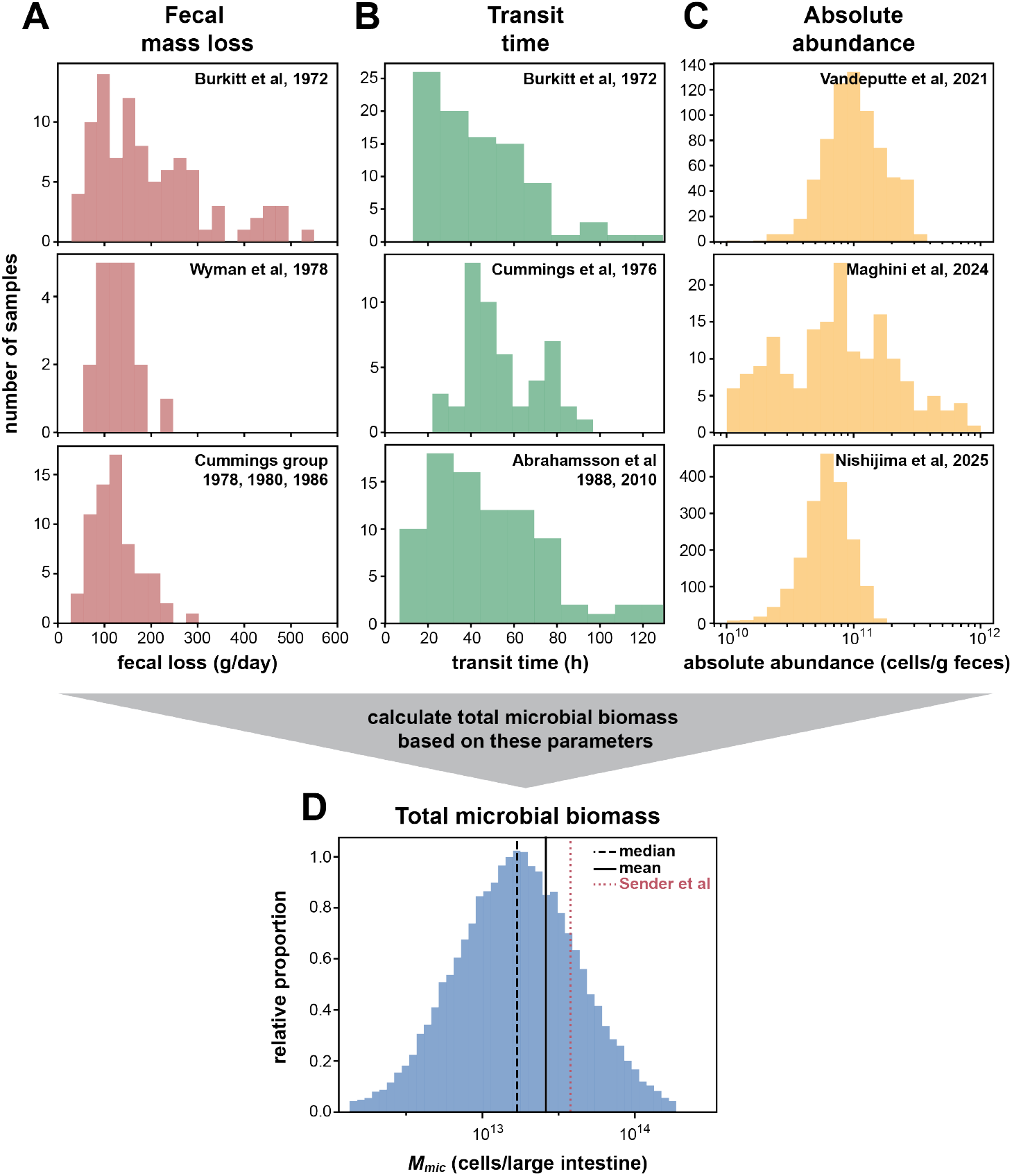
Empirical data on the variation in fecal mass loss, gastrointestinal transit time, and absolute bacterial abundance in feces. Representative datasets on (A) fecal mass loss [22, 25, 71–73], (B) gastrointestinal transit time [25, 70, 74, 75], and (C) absolute abundance in feces [15, 17, 21], show a strong variation of these quantities. (D) Proxy distribution of total bacterial biomass (bacterial cell counts) in the large intestine obtained by evaluating the mass-balance relation (Box 1) for different combinations of published measurements of fecal mass loss, gastrointestinal transit time, and absolute bacterial abundance. Mean *±* SD values are 170 *±* 100 g/day for fecal mass loss, 50 *±* 25 h for transit time, and 0.8 *×* 10^11^ *±* 1.1 *×* 10^11^ cells/g for absolute abundance (based on *n* = 180, 228, and 2806 measurements, respectively). Note that measurement techniques and study details, including participant selection, diet, and geographical region, vary across these studies (SI Text 2). Furthermore, the three variables are not independent. Thus, the distribution shown in (D) should be regarded as an illustrative proxy rather than the true population distribution. To generate this proxy, we combined the distributions of fecal mass loss and transit time with published distributions of absolute abundance, excluding the broad distribution reported by Maghini et al. (2024). Similar calculations using different subsets of the published datasets are shown in Figure S1.

Directly measuring these fluxes remains challenging. Chemical concentrations within the gut are difficult to obtain experimentally and provide only snapshots of a highly dynamic system with steep spatial and temporal gradients [8–10]. Here we introduce a systems-level analysis to estimate chemical flux magnitudes in the gut based on directly measurable quantities. In the following sections, we outline the logic underlying this analysis and discuss three major examples of chemical exchange: fermentation products, bacterial toxins, and nitrogen-containing metabolites. Our analysis provides explicit design guidance for future mechanistic studies to uncover health-relevant host-microbiome interactions.

## 2 Estimating the magnitude of chemical fluxes in the gut

To estimate the flux of chemicals exchanged between the microbiota and the host, we focus on the large intestine, the intestinal region that harbors the densest microbial community in the human body. We consider the mass balance of a microbially produced chemical *C* in the large intestine. This quantity is jointly determined by microbial production, host absorption, microbial degradation, fecal loss, and other possible processes (Figure 1B). The amount of the chemical will change over time according to the comparative magnitudes of these processes and their variation throughout the day. This can be formalized in a differential equation linking the temporal change in mass *C* to the magnitude of different flux terms (*J*_*i*_ in Figure 1C, Eq. 1). Importantly, when averaged over time scales long enough to account for diurnal variation in food intake and digestion, these complex relations simplify substantially, and the average host absorption per time period can be inferred from the mean rates of microbial production, microbial degradation and fecal loss. Mathematically, this corresponds to a quasi-steady state in which the fluctuations in the mass of the chemical *C* are averaged out over long time scales (Figure 1C, Eq. 2). In the simplest case, if the loss of *C* in feces and microbial degradation are negligible, the average host absorption flux is equivalent to the average microbial production flux, an approximation which holds well for fermentation products, as we discuss below. In any case, microbial production fluxes are the crucial starting point to estimate fluxes to the host.

So how can we estimate microbial production? Say a microbial species *S* is producing a chemical of interest, *C*. Then the production flux *J*_*C, pro*_ depends on the total biomass of this species in the large intestine *M*_*mic,S*_ and the rate *α* with which this species produces the chemical, *J*_*C, pro*_ = *αM*_*mic,S*_. While *α* can often be approximated through *in-vitro* measurements, as we discuss for specific examples below, we start with the determination of total microbial biomass *M*_*S*_ in the large intestine. This quantity is hard to measure directly, but as we demonstrate in the following, it can be estimated surprisingly well through a combination of accessible quantities starting with the abundance of bacteria in feces: Commonly, microbiome studies utilize DNA sequencing to report the *relative abundance* of microbial species in fecal samples (Table 1. i)[11]. More recently, it has been emphasized that going beyond compositional data to measure *absolute abundance* in microbiome samples is an important step forward [12, 13], and different approaches were introduced to measure or estimate the cell count or biomass of a bacterial species in fecal samples (Table 1. ii) [12–21]. These numbers describe bacterial abundance per sample weight, *A*_*abs,S*_. Themselves, these numbers are not sufficient to extrapolate the total microbial biomass in the large intestine. However, we can estimate this host-contextual quantity by incorporating measurements of fecal weight produced over time, the *fecal mass loss M*_*fec*_, and gastrointestinal transit time *t*_*transit*_: *M*_*mic,S*_ *≈ A*_*abs,S*_ *× M*_*fec*_ *× t*_*transit*_. This relation follows from bioreactor theory and robustly holds for the microbial growth and transport dynamics along the large intestine as we describe in detail in Box 1 and SI Text 1. In sum, combining absolute abundance measurements with two additional host-contextual quantities (transit time and fecal wet weight; see Table 1. iii & iv) allows the estimation of total microbial biomass or the biomass of a specific species *M*_*S*_ in the large intestine.

**Table 1:** The estimation of microbial biomass from abundance metrics, transit time, and fecal mass loss. Microbial biomass in the large intestine can be well estimated but requires, in addition to numbers on absolute abundance in feces, the measurement of two host-contextual quantities, total fecal mass loss and transit time, which cannot be derived from the analysis of fecal samples alone. All the quantities mentioned in the table can be measured using established methods.

|  | Quantity | Synonyms | Mathematical Relation | Units | Required information | Methods |  |
| --- | --- | --- | --- | --- | --- | --- | --- |
| i. | relative abundance |  |  | %, fraction | composition | sequencing | feces-derived metrics<br>(sample intrinsic) |
| | per species | rel. microbiome profile (RMP) | $A_{rel, S}$ | | | | |
| ii. | absolute abundance | | | g g<sup>-1</sup>, cells g<sup>-1</sup> | +<br>total abundance ($A_{abs}$) | qPCR, ddPCR, flow cytometry, DNA spike-in | |
| | total | microbial load | $A_{abs}$ | | | | |
| | per species | quant. microbiome profile (QMP) | $A_{abs, S} = A_{rel, S} \cdot A_{abs}$ | | | | |
| iii. | microbial loss in feces | | | g day<sup>-1</sup>, cells day<sup>-1</sup> | +<br>daily fecal mass loss ($M_{fec}$) | weighing | system-level metrics<br>(host contextual) |
| | total | | $J_{mic} = A_{abs} \cdot M_{fec}$ | | | | |
| | per species | | $J_{mic, S} = A_{abs, S} \cdot M_{fec}$ | | | | |
| iv. | microbial mass in large intestine | | | g, cells | +<br>transit time ($t_{transit}$) | dyes, radio opaque markers | |
| | total | | $M_{mic} = J_{mic} \cdot t_{transit}$ | | | | |
| | per species | | $M_{mic, S} = J_{mic, S} \cdot t_{transit}$ | | | | |

All three quantities can be determined by established and minimally invasive methods. Separate data on fecal mass loss, transit time, and absolute abundance are available in the literature and show strong variation between individuals and over time [13, 22] (Figure 2A–C). Using these data, we obtain a proxy for the distribution of total bacterial counts *M*_mic_ in the large intestine across individuals (Figures 2D and S1). Notably, the published point estimate most commonly cited for this quantity [23] lies within the range predicted by our analysis (Figure 2D, red dashed line). In contrast to that estimate, however, we do not assume an average gastrointestinal volume, which can vary substantially between individuals [24] and is difficult to measure directly. Instead, based on the fundamental relation coupling the microbial content within the large intestine to the microbial loss in feces (Box 1), we dynamically estimate bacterial counts from transit time, fecal mass loss, and absolute abundance. As these quantities are not independent variables [25, 26] and we combine values from different studies, the resulting distribution in Figure 2D should be regarded as an illustration. To determine *M*_mic_ within an individual and estimate its true population distribution, these three quantities must be measured simultaneously over sufficiently long time scales.

**Figure 2:**
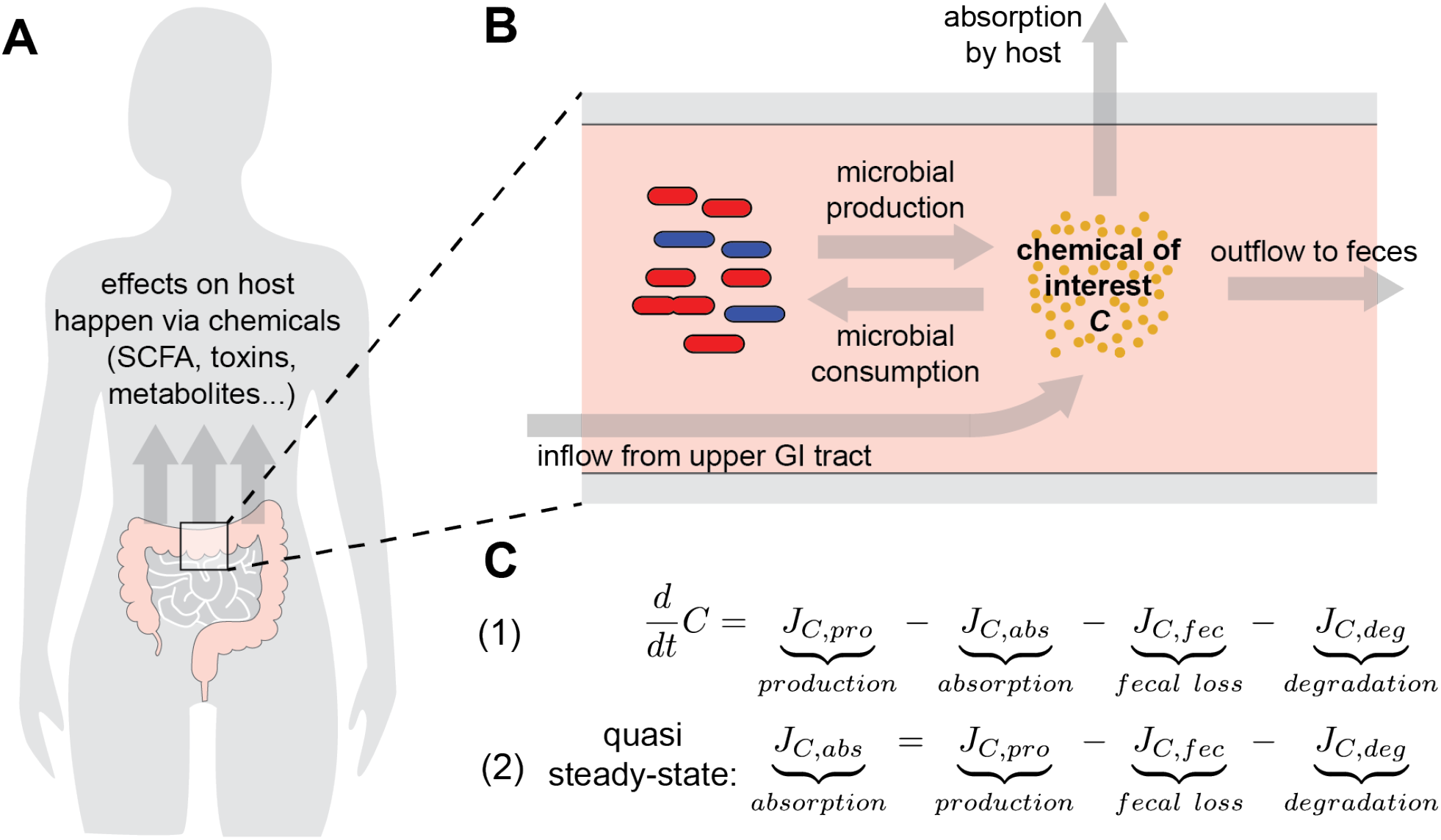
Balance of a microbially produced chemical in the large intestine. (A) The microbial community residing in the human large intestine affects host health and physiology via the exchange of various chemicals. (B) Schematic view of the different fluxes (arrows) contributing to the concentration of a chemical of interest in the gut lumen. (C) Equations that formally describe the balance of different fluxes. (1) The differential equation describing how the amounst of chemical *C* changes, depending on microbial production *J*_*C,pro*_, host absorption *J*_*C,upt*_, loss in feces *J*_*C, f ec*_, and degradation *J*_*C,deg*_. (2) Quasi-steady state equation of chemical *C*, for a timescale chosen to even out short-term fluctuations, 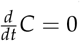. The absorption follows from production and loss via feces.

Finally, with an estimate of the total microbial mass *M*_mic_ and the respective mass of a single species *M*_mic,S_, we can return to our primary quantity of interest: the total production flux of a chemical of interest, *C*. As stated, this flux is obtained by combining *M*_mic,S_ with the microbial production rate per unit biomass, *α*, estimated, for example, from in vitro measurements, such that *J*_*C,pro*_ = *αM*_mic,S_. In the following, we illustrate this framework for three representative examples and show how it can be used to estimate the amount of microbial metabolites reaching host tissues.

## 3 The exchange of fermentation products

Bacterial fermentation products, including acetate, propionate, and butyrate, constitute the majority of chemicals exchanged between the gut microbiota and the host [26, 27]. They are the product of microbial energy metabolism within the anaerobic large intestine [28] and have well-documented roles in modulating host functions, such as immune regulation [29], epithelial cell metabolism [30], and satiety signaling [31]. While direct measurement of these fermentation fluxes is difficult, we recently estimated the total amount of microbiota-derived fermentation products reaching the host, using several orthogonal approaches[26]. Here, we recast this analysis within the general flux-estimation framework introduced above.

Three factors simplify the flux analysis in the case of fermentation products, leading to a very robust estimate of absorption by the host [26] (Figure 2A). First, fast-growing microbes, such as those in the gut [10, 32], require energy mostly for the synthesis of novel biomass (>90% of energy demand) [33, 34]. Thus, microbial fermentation is directly tied to the production of new microbial biomass. In our recent study, we have systematically quantified fermentation in 22 of the most abundant species in the human gut, showing that gut bacteria release fermentation products with an average rate per newly synthesized biomass of *α*_*FP*_ *≈* 29mmol (g bacterial dry weight)^*−*1^ [26]. Second, while directly measuring the synthesis of novel bacterial biomass in the large intestine is hard, this amount is close to the amount of microbes released in feces, *J*_*mic,growth*_ = *J*_*mic, f ec*_, as bacterial biomass lost in feces stems almost exclusively from growth in the large intestine [32, 35]. Therefore, the release of fermentation products by the gut microbiome follows as *J*_*FP,MB*_ *≈ α*_*FP*_ *· J*_*mic, f ec*_.

Third, the loss of fermentation products in feces is negligible [26]. We can thus estimate the host absorption of fermentation products directly from the release of fermentation products from the gut microbiota, *J*_*FP,absorp*_ *≈ J*_*FP,mic*_ *≈ α*_*FP*_ *· J*_*mic, f ec*_. As discussed above, *J*_*mic, f ec*_ can be determined using data on fecal mass loss and the absolute abundance of microbes in feces (Table 1, iii). Accordingly, the flux of fermentation products to the host can be estimated from directly measurable quantities.

This estimation provides highly consistent absorption numbers, as we have systematically analyzed, integrating metabolic measurements, the quantification of consumed complex carbohydrates, and a comparison between germ-free and conventional mice[26].

## 4 Toxin fluxes during *C. difficile* infections

As a second example we apply our framework to toxin fluxes. Toxins released by gastrointestinal pathogens such as *Clostridioides difficile, Vibrio cholerae*, and *Shigella* spp. can cause severe diarrhea and other symptoms [36]. A better understanding of toxin fluxes is a key step towards a mechanistic and actionable understanding of the infection process: How much toxin does a pathogen produce in a given microbiota, how does toxin production and turnover depend on the physiological conditions along the gut, and how do these factors interact to set toxin levels reaching the host? Below, we illustrate this flux analysis using *C. difficile* as an example.

While the abundance of *C. difficile* is typically low in a healthy gut (below 0.01% relative abundance and ≪ 10^6^ cells per g feces), antibiotic treatment can lead to blooming *C. difficile* populations (around 0.1 to 10% of relative abundance, or 10^6^ to 10^9^ cells per g feces) [37–39]. The *C. difficile* toxins TcdA and TcdB disrupt epithelial integrity and cause diarrhea [7], with disease severity correlating with fecal toxin levels [40]. Therefore, knowledge of the amount of toxin produced by *C. difficile* and the emerging toxin concentration in the gut is a critical step to understand the onset and progression of the infection. To analyze this toxin flux, we first consider the toxin release by *C. difficile*. In-vitro studies have shown that toxin release varies broadly with environmental conditions and between strains. To approximate relevant levels of toxin production without specific knowledge of gene regulation in a specific microbiota, we combine data on time-resolved toxin concentrations and bacterial biomass [41–43], and find production rates per bacterial dry weight *α*_*tox*_ to fall within a range of 0.1 and 1 *ng µg*^*−*1^ *h*^*−*1^. The total production flux of toxins, *J*_*tox,pro*_, depends on this rate and the biomass *M*_*Cdi f f*_ of *C. difficile* in the gut, *J*_*tox,pro*_ = *α*_*tox*_ *M*_*Cdi f f*_. The total mass of toxins *T* in the gut in turn depends on this production flux and the loss via feces (Figure 2B). Following bioreactor theory, the loss can be approximated by *J*_*tox, f ec*_ = *T*/*t*_*transit*_; as content moves through the gut faster and transit times are shorter, more toxins are lost via feces. Averaged over time and converting to concentrations (SI Text 3), the toxin concentration [*T*] follows as

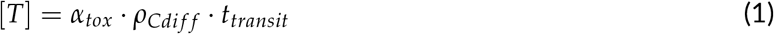

Here, *ρ*_*Cdi f f*_ is the density of *C. difficile* in gut content which can be estimated from the absolute abundance of C. difficile and the mass density of microbes in feces (SI Text 3).

With this relation, we can estimate toxin concentrations for an acute infection. Typical densities of *C. difficile* in stool are around *ρ*_*Cdi f f*_ = 1 mg dry mass per mL feces [40, 44–47]. As a secretory diarrhea, severe *C. difficile* colitis is associated with a markedly shortened gastrointestinal transit time, typically falling below 12 hours during acute disease [48]. Under these conditions, estimated in situ toxin concentrations reach around 1*µg*/*mL*, comparable to the upper range of toxin concentrations measured in stool during acute infections [49].

This simple flux consideration highlights the dual role of diarrhea as a systemic response to infection. First, diarrhea and faster transit times increase the loss of toxins via feces, reducing toxin concentrations. Second, diarrhea and faster transit times increase the loss of *C. difficile* itself, further reducing the synthesis of new toxins. For a more mechanistic understanding of *C. difficile* disease progression, it will be essential to routinely measure transit times and fecal toxin concentrations during acute infections, and to systematically investigate toxin production rates *α*_*tox*_ across intestinally relevant conditions and *C. difficile* strains.

## 5 The role of the gut microbiota in nitrogen homeostasis

As a third example, we consider nitrogen fluxes in the gut. Microbes can degrade and utilize many of the major nitrogen containing compounds in the gut [50–52] (Figure 2C), with strong impacts on nitrogen homeostasis and the health of the host. For example, microbial ammonia production has been linked to hyperammonemia and hepatic encephalopathy [53, 54], and the microbial degradation of peptides can lead to the production of toxic byproducts such as hydrogen sulfide [55]. In addition, bacterial growth varies with the availability of nitrogen sources [56] with potential consequences on microbiome composition. Despite this relevance, the quantitative contributions of different nitrogen-modifying processes to the gut’s nitrogen balance are not well mapped out. Using the framework of flux estimation provides a good starting point.

As an example, we discuss specifically ammonia fluxes in the gut. Several processes affect the ammonia (*NH*_3_) pool in the gut (Figure 2C ii). It can be produced by microbial activity from protein, urea, or mucin [50]. *NH*_3_ also enters the large intestine from the upper gastrointestinal tract [57]. *NH*_3_ is the preferred nitrogen source for highly abundant gut bacteria [58] which can sequester it in bacterial biomass. *NH*_3_ can also enter the blood stream of the host or be lost via feces.

To understand the turnover of *NH*_3_ and thus the amount reaching the host, we have to estimate the magnitudes of these different processes. Averaged over time, the *NH*_3_ flux to the host depends on the magnitude of the other fluxes (Figure 2C iii). We can put numbers on some of the flux terms in this equation. For example, it has been estimated that bacteria break down a total of around 6g urea per day [50], that around 7g of protein arrive in the large intestine undigested [26], and that 4.4g of mucin is available for bacterial digestion in the large intestine[26]. When it comes to microbial activity, much less is known. There is an increasing understanding of the nitrogen transforming capacity of different microbiota members. For example, the ability of microbes to break down urea has recently been mapped out and functionally assessed [59–62]. However, the activities of the involved metabolic pathways and how their throughput is shaped by the environmental conditions bacteria encounter in the gut is still largely unclear.

The flux centric view of intestinal *NH*_3_ turnover underscores the need to better understand the quantitative contribution of different microbial groups to *NH*_3_ production and sequestration into biomass. *NH*_3_ fluxes in the gut can be estimated based on the *NH*_3_ consumption and production rates and the total biomass of bacteria performing these functions. Importantly, while we use *NH*_3_ summarily for both, ammonia and ammonium 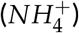 in this section, those molecules are in a pH-dependent equilibrium, and have different properties with regard to solubility and bioavailabilty. Rates of biologically relevant reactions can be estimated through systematic in-vitro studies across microbiota members, while total bacterial biomass follows from absolute abundance in fecal samples, transit time, and fecal loss measurements as discussed before. Overall, this highlights the future potential of flux estimations to quantitatively understand the role of gut bacteria in the host’s nitrogen homeostasis and its impact on health.

## 6 Discussion

Much like pharmaceutical or environmental agents act in a dose-dependent manner, the impact of the human microbiome on host physiology depends on the magnitude of the chemical fluxes it exchanges with the host. Quantifying these fluxes is therefore essential to understand how the microbiome shapes health and disease. Here, we analyzed the host-microbiota system from first principles to identify the core determinants of these fluxes. Microbiota functions depend on the abundance and activity of bacteria in the system. Both of these quantities are closely related to growth and turnover of bacterial biomass in the large intestine. Connecting these processes through bioreactor theory, we identify the major determinants of chemical fluxes and how they relate to each other.

These relations highlight three host-level parameters that, together with microbial activity, shape flux magnitudes: absolute microbial abundance in feces, gastrointestinal transit time, and fecal loss. Notably, although these quantities have rarely been analyzed together, transit time and absolute abundance have been identified in recent microbiome studies as key determinants of microbiome composition and microbe–host interactions. Absolute microbial abundance in fecal samples has been repeatedly found to correlate strongly with microbiome composition and diversity, characteristics of host immunology and digestion, and a range of disease symptoms[13, 15, 20, 21]. Transit time has been shown to relate to the metabolic activity of the microbiome and the metabolic footprint it leaves in the blood [63, 64]. It is also strongly correlated with stool energy density and the composition and diversity of the microbiome [65–68]. Given that absolute abundance, transit time, and fecal loss are major determinants of chemical fluxes in the gut, these three quantities should routinely be measured together. While transit time and absolute abundance have received considerable attention in recent microbiome studies, fecal mass loss data remain largely absent from modern work despite being relatively straightforward to assess. We therefore specifically call for the systematic inclusion of fecal mass loss measurements in future studies.

In addition we need to understand how microbial activity shapes chemical fluxes. While absolute abundance, transit time, and fecal loss together determine the total biomass of microbes in the gut, we also need to estimate the per-biomass rates at which these microbes produce or consume a chemical of interest. These rates arise from microbial physiology and metabolism, which differ across microbial species and vary with environmental conditions, such as the type and availability of nutrients. Systematic in vitro experiments, which can yield per-biomass transformation rates of a given chemical across strains and gut-relevant conditions, are crucial for understanding microbial activity and predicting chemical fluxes.

In conclusion, by incorporating a set of measurable quantities, the reasoning introduced here provides a feasible framework for estimating chemical fluxes between the gut microbiota and the host. This provides a missing input for studying pharmacokinetics and pharmacodynamics of microbiota-derived compounds, critical to dissect their effects in the human body. When applied systematically across different individuals, diverse diets, and varied microbiomes, our framework can enable mechanistic insights into the molecular exchanges that underlie microbiome–host interactions in health and disease.

## Acknowledgments

We thank members of the Cremer group, Emma Slack and Sebastian Hummel for critical reading and helpful comments. JC acknowledges support from Food@Stanford. MA was supported as part of NCCR Microbiomes, a National Center of Competence in Research, funded by the Swiss National Science Foundation (grant number 180575).

## Box 1

**Estimating microbial biomass in the human large intestine**

The vast majority of gut microbes grow in the proximal large intestine and eventually exit the intestinal tract via feces [10].These dynamics establish a fundamental relation between fecal loss, gastrointestinal transit time, and the total microbial biomass present in the large intestine. To illustrate this, we consider a well-mixed bioreactor with constant microbial cell density *ρ* as an example (Figure). The total mass of microbes in the reactor *M*_*mic*_ follows from the density *ρ* and the total volume *V* of the reactor, *M*_*mic*_ = *ρV*. The rate of bacterial biomass loss via outflow, *J*_*mic, out*_, scales with the density *ρ* and the flow rate *Q*_*V*_ quantifying how much volume is passing through the reactor per time, *J*_*mic, out*_ = *ρQ*_*V*_. If we divide both quantities, we get

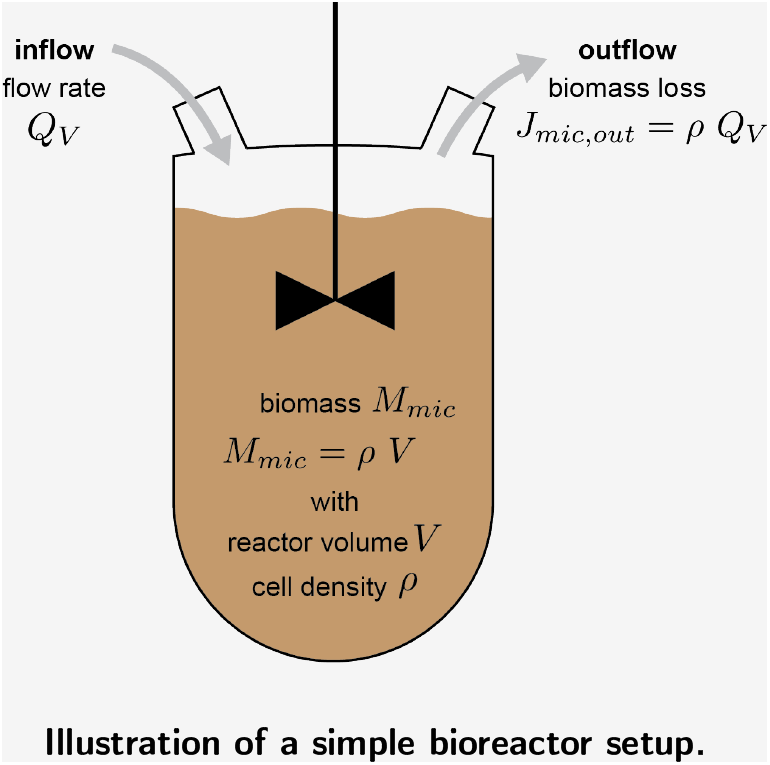

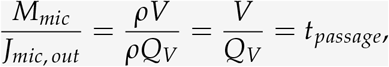

where the ratio *V*/*Q*_*V*_ of volume to outflow rate is the time it takes to exchange the volume of the bioreactor, or the average passage time *t*_*passage*_. Thus, the total microbial mass in the bioreactor is related to the loss of biomass by

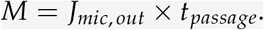

For the large intestine, *t*_*passage*_ can be well approximated by the total gastrointestinal transit time *t*_transit_ because gastric and small-intestinal transit are relatively fast in comparison: meals empty from the stomach with a half time of one to two hours and reach the colon after approximately three hours [69] whereas whole-gut transit times commonly range from 12 to 60 hours [70] (Figure 2, central column). The average rate of microbial biomass loss in feces *J*_*mic,out*_ follows from the absolute abundance of microbes in feces *A*_*abs*_ multiplied by the fecal mass loss *M*_*f ec*_. While *M*_*f ec*_ can vary strongly with discrete defecation events over short timescales, these events average out when measured over longer time scales, ideally several days. The microbial biomass in the large intestine can then be estimated as:

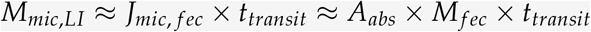

Importantly, this relation is not restricted to well-mixed conditions; it also applies to more realistic flow regimes, including plug-flow-reactor models where microbial growth occurs near the inlet and microbial biomass moves with the flow along the tube (SI Text 1). Furthermore, the transit time *t*_*transit*_, the absolute abundance in feces *A*_*abs*_ and daily fecal mass loss *M*_*f ec*_ are host-contextual quantities that can be directly measured (see main text). To estimate the biomass of specific species, we can further integrate abundance data from sequencing studies (Table 1 iv).

## Figures and Tables

**Figure S1:**
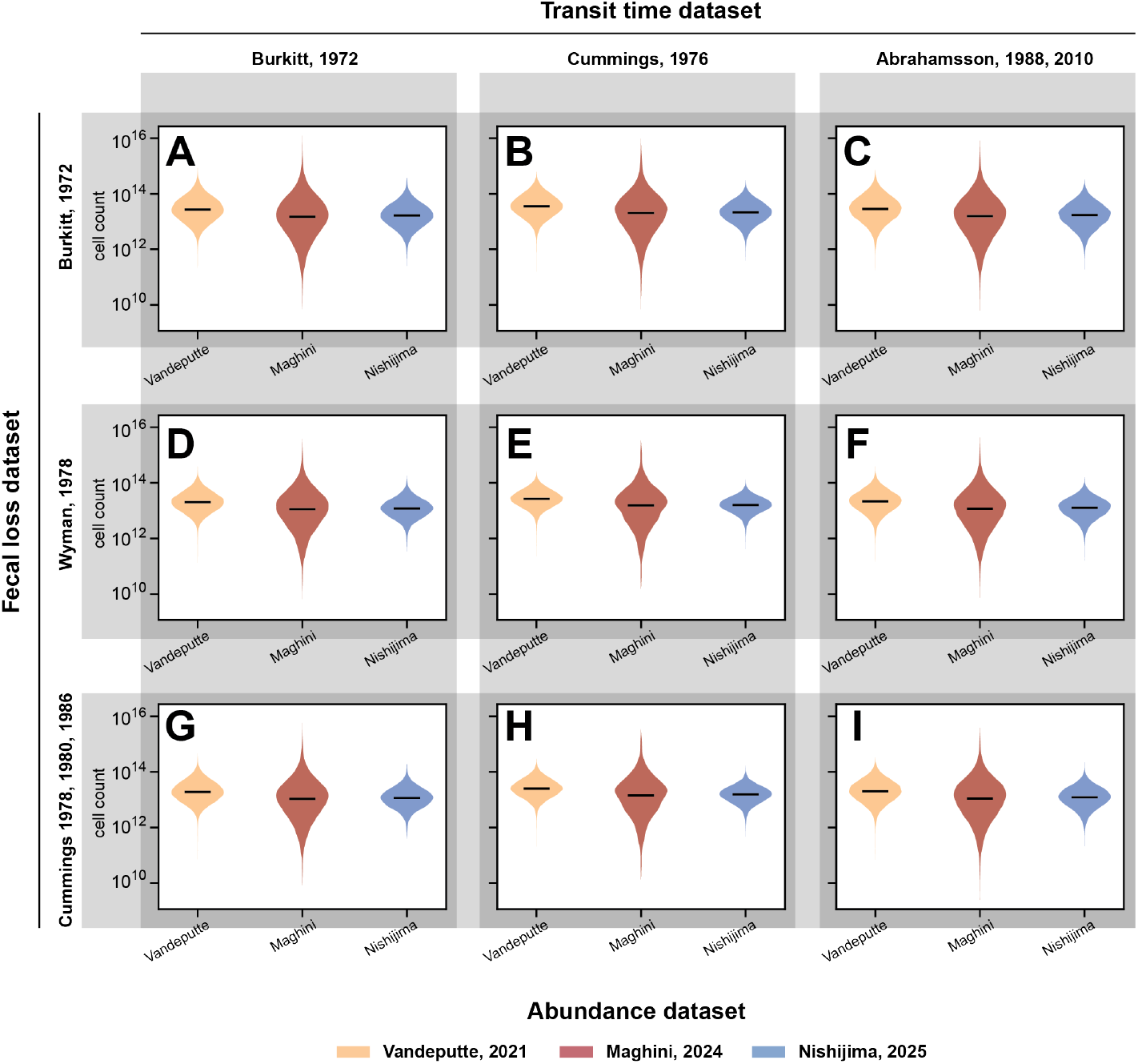
Estimating variation in total large intestinal cell count based on published data on the variation in fecal mass loss [22, 25, 71–73], gastrointestinal transit time [25, 70, 74, 75], and absolute bacterial abundance [15, 17, 21]. Every row of plots shows data based on a different dataset for fecal mass loss, while every column of plots shows data based on a different dataset for gastrointestinal transit times. Each plot has data for three different datasets for total bacterial abundance in feces.

## Supplementary Information

### 1 Estimating bacterial abundance in flow reactors and the large intestine

Bioreactor theory describes how bacterial biomass in systems with turnover depends on nutrient supply, microbial growth, flow conditions, and the rate of biomass loss from the system. It has been applied to the intestines of different animals with diverse anatomy and physiology to better understand how microbial biomass in distinct intestinal sections, bacterial loss, and gastrointestinal transit times are inherently linked [1–3]. Historically, this approach has been most extensively developed and applied to understand microbial biomass turnover in ruminants, but it is increasingly being used to obtain more detailed models of microbial growth and biomass turnover in the human gut [4–10].

Growth and biomass turnover in the human large intestine are highly dynamic, with strong diurnal variations in growth, fluid flow, and biomass turnover. However, the total microbial mass *M*_*mic,LI*_ in the human large intestine still follows, to first order, from the loss of microbes in feces *J*_*mic, f eces*_ and the gastrointestinal transit time *t*_*transit*_:

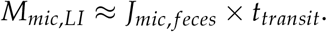

In the main text, we considered a well-mixed bioreactor with nutrient inflow and biomass loss to introduce this relation. The human large intestine differs substantially from this simple scenario, exhibiting a tube-like geometry without strong mixing, pronounced diurnal variation in flow, and active water uptake along the colon, in addition to nutrient inflow and fecal loss. Here, we discuss why the fundamental relation between microbial mass, gastrointestinal transit time, and fecal loss nevertheless holds robustly beyond the well-mixed case.

First, the relation also holds for tube-like geometries. For illustration, consider a tube of length *L* containing microbes with constant density *ρ* moving with velocity *v* through the tube. In such a plug-flow reactor it takes *t*_*passage*_ = *L*/*v* for biomass to traverse the system, and the total biomass in the tube follows from the biomass loss rate as *M*_*mic*_ = *J*_*mic, f eces*_ *× t*_*passage*_, identical to the relation derived for the well-mixed scenario described in the main text. This simple plug-flow-reactor scenario assumes a constant bacterial density. This holds approximately for the large intestine: The adult human large intestine is approximately 150–180 cm long. Supplied by complex carbohydrates and other nutrients entering from the small intestine, microbial growth primarily occurs in the proximal region of the large intestine, the cecum and ascending colon. In contrast, microbial density in the distal large intestine (dLI) beyond the ascending colon is more constant. As the cecum and ascending colon are only about 20 to 30 cm long and exhibit lower microbial densities than the distal parts of the large intestine, the majority of microbial biomass resides in the more distal regions with more constant densities. As such, the mass balance relation holds well:

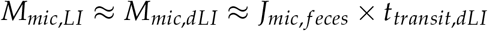

Beyond these considerations, water absorption by the host can also alter microbial densities, potentially modifying the mass balance equation of the plug-flow reactor. However, most water absorption occurs in the proximal colon, making it again less relevant in modifying microbial densities along the distal parts of the gut. Furthermore, by removing a large fraction of the water entering from the small intestine, commonly 85 to 90%, water absorption also reduces the flow velocity of luminal content along the large intestine. As a consequence, transit time is also largely determined by passage through the more distal regions, after most water uptake has already occurred, *t*_*transit*_ *≈ t*_*transit,LI*_ *≈ t*_*transit,dLI*_

Taken together, the relation *M*_*mic,LI*_ *≈ J*_*mic, f ec*_ *× t*_*transit*_ reflects a fundamental interdependence between biomass, transit time, and fecal loss, providing a robust and practical estimate of microbial biomass in the human large intestine.

### 2 Quantification and datasets of fecal loss, transit time, and absolute abundance

To illustrate inter-person variation in fecal loss, intestinal transit time, and the absolute abundance of microbes in feces, we show in Figure 3 three major datasets from the literature for each of these quantities. Here, we introduce study conditions and methods involved. We also discuss the correlations between the three quantities.

**Figure 3:**
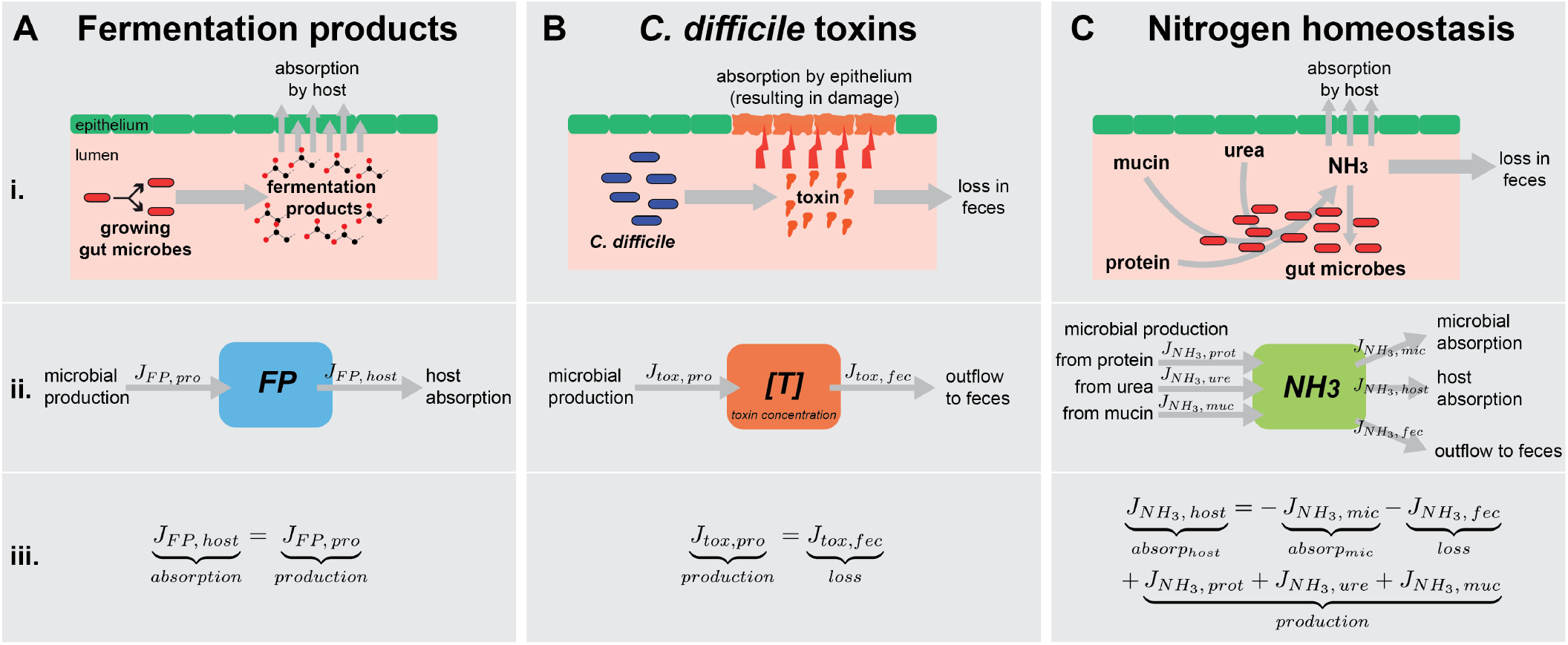
Three examples for flux estimations in the gut microbiota. (A) Microbiota-derived fermentation products. As in the large intestine, both inflow of fermentation products from the upper gastrointestinal tract, as well as loss of these molecules via feces are negligible, host absorption is largely determined by microbial production. (B) *C. difficile* toxin concentrations. Concentration resulted from the balance of toxin production and the loss via feces. (C) Nitrogen homeostasis in the gut. Ammonium (NH3) turnover is the combined result of several processes requiring different flux estimations.

### 2.1 Fecal mass loss

Weighing fecal mass loss is conceptually simple, and, as we argue in the main text, very important to understand chemical fluxes in the gut. Nevertheless, fecal loss is rarely measured in modern microbiome studies. The comprehensive datasets we chose to show in Figure 3 are from older studies, dating back more than 30 years.

1. Burkitt et al, Lancet 1972. This dataset is the most comprehensive one on fecal loss. It is derived from a seminal study by Burkitt and colleagues comparing fecal loss and transit time in 94 volunteers that have different backgrounds, both in terms of wether they were living in more rural or urban areas, and whether they consumed a more westernized or more traditional diet. Geographically, the study has a focus on Africa and Europe. Notably, as the data was collected around 1970, even the western diet was more rich in fiber and contained less sugars [11]. The study participants were asked to collect five or six consecutive stools in plastic bags, and the weight of these stools was then used to assess daily fecal loss.
2. Wyman et al, Gut 1978. The goal of this study was to understand variation in the amount and consistency of passed stool between people and in the same person over time. They recruited 20 volunteers (10 male, 10 female), most of which were adults (24-48 years old), except for one boy (11) and one girl (15). The men were given dietary instructions aimed at standardizing their fiber intake, while the women continued eating their normal diet, but were asked to consume the same amount of fiber-containing foods. All participants were provided kits for stool collection, and collected their stools for five consecutive days.
3. For the third example, we pooled data from three studies conducted by the same group, which used very similar methods. (i) Cummings et al, Lancet, 1978. This study was designed to test high-fiber interventions, and was performed on 19 male adult volunteers. We used only the data on the control diet (no increased dietary fiber content) in Figure 3, which was obtained through stool collections for three weeks, while the participants were provided a controlled basal diet which was meant to reflect a normal British diet at the time. (ii) Stephen and Cummings, J. Med. Microbiol, 1980. Aimed at measuring the microbial contribution to fecal output, this study used data from 9 male volunteers, who collected feces for three weeks while consuming a controlled diet reflecting standard British-type nutrition. (iii) Stephen et al, British Journal of Nutrition, 1986. 30 volunteers were recruited to test the effect of age, sex, and fiber intake on colonic function. The subjects were provided four different diets for a three week period, and feces were collected on days 13-21 of this time.

### 2.2 Gastrointestinal transit time

Different methods have been developed to estimate gastrointestinal transit times and transit times through specific intestinal sections. The typically rely on the ingestion of radio-opaque markers, which can then be visualized using repeated abdominal X-ray imaging, X-ray imaging of consecutively passed feces, or a single abdominal X-ray after continuous marker ingestion over sufficiently long times to allow quasi-steady state marker counts. Here, we chose datasets which systematically quantified transit times and report transit times in units of time, and opted to show studies that have introduced new methods, together with more recent work.

1. Burkitt et al, Lancet 1972. This the same study as for the fecal loss described above. Transit time was measured by ingestion of 25 radio-opaque plastic pellets. The next five to six stools were collected and the times of collection were noted. The stools were then analyzed using X-ray imaging, and the time when 20 of the pellets were excreted was calculated and called the transit time.
2. Cummings et al 1976: This is the first study where steady state marker counts using continuous marker intake were used, and the authors compare this method with recovery in stool of markers from a single dose over time. Twelve male subjects participated in several consecutive study protocols, including continuous and single marker administration, and different diets (ad libitum, controlled western style, high fiber). In Figure 2, we use mean transit time data derived from 50 continuous marker studies across diets.
3. For the third example, we used two studies from the same researchers using the same method, but published 22 years apart. (i) Abrahamsson et al, Scand. J. Gastroenterol 1988. Uses continuous intake of radioopaque markers for three days followed by one abdominal X-ray. 23 healthy adult men and 33 healthy adult women were studied. They kept their usual habits with regard to eating, drinking, smoking, and physical activity, but recorded these factors. (ii) Abrahamsson et al, Neurogastroenterol Motil 2010. Here, the goal was testing how many markers need to be taken to attain a steady state marker presence in people with different transit times. Fifteen healthy people (ten men, five women), and thirteen people (four males, nine females) with altered bowel habits were studied, following normal dietary and lifestyle habits. The received different numbers of markers for different periods of time. We used data generated using 50 ingested markers.

### 2.3 Absolute abundance

Methods to estimate the total abundance of bacteria in fecal samples either rely on methods that quantify the number of cells directly, mostly through flow cytometery, or the estimation of cell numbers using sequence-based methods (qPCR, sequencing with normalization). In Figure 2, we use two datasets that use flow cytometry, and one using qPCR.

1. Vandeputte et al, Nature Communications, 2021. In this study, the authors collected fecal time series of 20 healthy adult women (age 16-55), 713 samples in total, to assess the inter and intra individual variation in species composition and absolute abundance.
2. Nishijima et al, Cell 2025. This study uses 1894 available samples from the study cohorts GALAXY / MicrobLiver and MetaCardis [12–14], and uses flow cytometry-based quantification.
3. Maghini et al, Nature Biotechnology, 2024. This dataset if based on 16S qPCR. The study includes 10 adult donors that provided a single stool sample, with the goal to evaluate storage temperature and choice of preservative for measurements of microbial load and composition. With all combinations of preservation methods, ever sample was analyzed 21 times, giving a total of 210 measurements. The study reports absolute abundance per gram dry weight, while the others report it per total fecal weight. To enable comparison, we used a factor of 0.3g dry weight per g wet weight to renormalize the data.

For analyzing the data reported in the datasets above, we used numerical data reported in the original studies when available. In cases where data was only reported in plots, we used an online tool (WebPlot-Digitizer, https://github.com/automeris-io/WebPlotDigitizer) to extract numerical data from the plots.

### 2.4 Correlation between quantities

It is important to note that fecal mass loss, gastrointestinal transit time, and absolute abundance might correlate. If there was a strong correlation, it could even be possible to infer one quantity from measurements of the other. Fecal mass loss and transit time have historically often been measured together. The data from Burkitt et al mentioned above is a good example. However, in that study there is only an apparent correlation at very low values for fecal mass loss (below around 120g/day), where transit time decreases strongly if fecal mass increases. This relationship is much weaker at higher values for fecal mass loss, where transit time seems to vary little (Figure S9 in [11]).

Finding good data to predict correlations with absolute abundance is harder. While Nishijima report that absolute abundance co-varies with many host parameters, the association with transit time (with Bristol Stool Scale measurements as a proxy) is not very strong, and fecal mass loss measurements are not available in these cohorts.

Overall, it is thus important to keep in mind that the three variables are not necessarily independent. However, correlations are by no means certain and all three quantities should still be measured simultaneously to promote better flux estimations.

### 3 Concentration conversions for *C. difficile* toxins and other chemicals

In the maintext, we introduced flux estimations in terms of absolute numbers and total fluxes in the host. That is, we estimated the total amount of a chemical within the large intestine, depending on the different fluxes modifying this chemical. However, it might often be more relevant to consider chemical concentrations instead of absolute amounts.

Depending on production, absorption, modification, and fecal loss of a chemical, what is its concentration in the large intestine? This should especially be true for bacterial toxins, such as those produced by *C. difficile* we consider in the main text. For their impact on epithelial health, understanding their concentration at the epithelium barrier is more important than knowing the total amount. Here, we describe how our flux considerations can be converted from total amounts to concentrations.

Concentrations are defined by the amount of a chemical *C* per volume *V*, [*C*] = *C*/*V*. The concentration of a chemical in the large intestine follows from the absolute abundance of this chemical in the large intestine divided by the volume of the intestinal lumen. While both volumes and absolute abundance measurements are hard to obtain, we can approximate the concentration in the large intestine by the concentration in feces, assuming little variation in concentrations throughout most of the large intestine.

For example, consider the *C. difficile* toxin in the large intestine. Production and loss via feces are given by *J*_*tox,pro*_ = *α*_*tox*_ *M*_*Cdi f f*_ and *J*_*tox, f ec*_ = *T*/*t*_*transit*_, and the total amount of toxin in the large intestine is given by *T* = *α*_*tox*_ *M*_*Cdi f f*_ *t*_*transit*_. To approximate the concentration of toxin [*T*] we can divide the total amount *T* by the volume of feces and calculate:

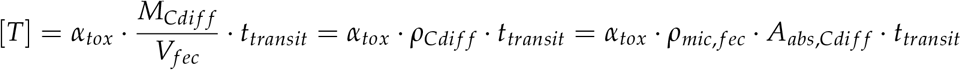

The density of *C. difficile* in feces (*ρ*_*Cdi f f*_, e.g. in units *g*/*mL*) can be approximated by the total density of bacteria (*ρ*_*mic, f ec*_ e.g. in units *g*/*mL*) and the absolute abundance of *C. difficile* in feces (*A*_*abs,Cdi f f*_ e.g. in units g/g).

## References

1. Marchesi, J. R. et al. The Gut Microbiota and Host Health: A New Clinical Frontier. Gut 65, 330–339. ISSN: 0017-5749, 1468-3288. (2025) (Feb. 2016).

2. De Vos, W. M., Tilg, H., Van Hul, M. & Cani, P. D. Gut Microbiome and Health: Mechanistic Insights. Gut 71, 1020–1032. ISSN: 0017-5749, 1468-3288. (2025) (May 2022).

3. Koh, A., Vadder, F. D., Kovatcheva-Datchary, P. & Bäckhed, F. From Dietary Fiber to Host Physiology: Short-Chain Fatty Acids as Key Bacterial Metabolites. Cell 165, 1332–1345 (June 2016).

4. Yano, J. M. et al. Indigenous Bacteria from the Gut Microbiota Regulate Host Serotonin Biosynthesis. Cell 161, 264–276 (Apr. 2015).

5. Otaru, N. et al. GABA Production by Human Intestinal Bacteroides Spp.: Prevalence, Regulation, and Role in Acid Stress Tolerance. Frontiers in Microbiology 12, 656895. ISSN: 1664-302X. (2022) (Apr. 2021).

6. Tarracchini, C. et al. Exploring the Vitamin Biosynthesis Landscape of the Human Gut Microbiota. mSystems 9 (ed Gilbert, J. A.) e00929–24. ISSN: 2379-5077. (2025) (Oct. 2024).

7. Aktories, K., Schwan, C. & Jank, T. Clostridium Difficile Toxin Biology. Annual Review of Microbiology 71, 281–307. ISSN: 0066-4227, 1545-3251. (2025) (Sept. 2017).

8. Phillips, S. F. & Giller, J. The Contribution of the Colon to Electrolyte and Water Conservation in Man. The Journal of Laboratory and Clinical Medicine 81, 733–746. ISSN: 0022-2143 (May 1973).

9. Hoces, D. et al. Metabolic Reconstitution of Germ-Free Mice by a Gnotobiotic Microbiota Varies over the Circadian Cycle. PLOS Biology 20 (ed Suez, J.) e3001743. ISSN: 1545-7885. (2022) (Sept. 2022).

10. Salari, A. & Cremer, J. Diurnal Variations in Digestion and Flow Drive Microbial Dynamics in the Gut. PRX Life 3, 023012. ISSN: 2835-8279. (2025) (June 2025).

11. Gloor, G. B., Macklaim, J. M., Pawlowsky-Glahn, V. & Egozcue, J. J. Microbiome Datasets Are Compositional: And This Is Not Optional. Frontiers in Microbiology 8. ISSN: 1664-302X. (2025) (Nov. 2017).

12. Morton, J. T. et al. Establishing Microbial Composition Measurement Standards with Reference Frames. Nature Communications 10. ISSN: 2041-1723. (2025) (June 2019).

13. Vandeputte, D. et al. Quantitative Microbiome Profiling Links Gut Community Variation to Microbial Load. Nature 551, 507–511 (Nov. 2017).

14. Barlow, J. T., Bogatyrev, S. R. & Ismagilov, R. F. A Quantitative Sequencing Framework for Absolute Abundance Measurements of Mucosal and Lumenal Microbial Communities. Nature Communications 11, 2590 (2020).

15. Vandeputte, D. et al. Temporal Variability in Quantitative Human Gut Microbiome Profiles and Implications for Clinical Research. Nature Communications 12. ISSN: 2041-1723. (2025) (Nov. 2021).

16. Jian, C., Luukkonen, P., Yki-Järvinen, H., Salonen, A. & Korpela, K. Quantitative PCR Provides a Simple and Accessible Method for Quantitative Microbiota Profiling. PLOS ONE 15 (ed Vaz-Moreira, I.) e0227285. ISSN: 1932-6203. (2025) (Jan. 2020).

17. Maghini, D. G. et al. Quantifying Bias Introduced by Sample Collection in Relative and Absolute Microbiome Measurements. Nature Biotechnology 42, 328–338. ISSN: 1087-0156, 1546-1696. (2025) (Feb. 2024).

18. Stämmler, F. et al. Adjusting Microbiome Profiles for Differences in Microbial Load by Spike-in Bacteria. Microbiome 4. ISSN: 2049-2618. (2025) (Dec. 2016).

19. Wirbel, J. et al. Accurate Prediction of Absolute Prokaryotic Abundance from DNA Concentration. Cell Reports Methods 5, 101030. ISSN: 2667-2375. (2025) (May 2025).

20. Contijoch, E. J. et al. Gut Microbiota Density Influences Host Physiology and Is Shaped by Host and Microbial Factors. eLife 8, 337–26. ISSN: 2050-084X (Jan. 2019).

21. Nishijima, S. et al. Fecal Microbial Load Is a Major Determinant of Gut Microbiome Variation and a Confounder for Disease Associations. Cell 188, 222–236.e15. ISSN: 0092-8674. (2025) (Jan. 2025).

22. Cummings, J. et al. Colonic Response to Dietary Fibre from Carrot, Cabbage, Apple, Bran, and Guar Gum. The Lancet 311, 5–9. ISSN: 01406736. (2021) (Jan. 1978).

23. Sender, R., Fuchs, S. & Milo, R. Revised Estimates for the Number of Human and Bacteria Cells in the Body. PLOS Biology 14, e1002533. ISSN: 1545-7885. (2024) (Aug. 2016).

24. Pritchard, S. E. et al. Fasting and Postprandial Volumes of the Undisturbed Colon: Normal Values and Changes in Diarrhea-Predominant Irritable Bowel Syndrome Measured Using Serial MRI. Neurogastroenterology & Motility 26, 124–130 (Oct. 2013).

25. Burkitt, D., Walker, A. & Painter, N. Effect of Dietary Fibre on Stools and Transit-Times, and Its Role in the Causation of Disease. The Lancet 300, 1408–1411. ISSN: 01406736. (2021) (Dec. 1972).

26. Arnoldini, M. et al. Quantifying the Varying Harvest of Fermentation Products from the Human Gut Microbiota. Cell 188, P5332–5342.E16. ISSN: 0092-8674. (2025) (Sept. 2025).

27. den Besten, G. et al. The Role of Short-Chain Fatty Acids in the Interplay between Diet, Gut Microbiota, and Host Energy Metabolism. Journal of Lipid Research 54, 2325–2340. ISSN: 0022-2275 (Aug. 2013).

28. Mukhopadhya, I. & Louis, P. Gut Microbiota-Derived Short-Chain Fatty Acids and Their Role in Human Health and Disease. Nature Reviews Microbiology 23, 635–651. ISSN: 1740-1526, 1740-1534. (2025) (Oct. 2025).

29. Corrêa-Oliveira, R., Fachi, J. L., Vieira, A., Sato, F. T. & Vinolo, M. A. R. Regulation of Immune Cell Function by Short-Chain Fatty Acids. Clinical & Translational Immunology 5, e73. ISSN: 2050-0068. (2021) (Apr. 2016).

30. Roediger, W. E. Role of Anaerobic Bacteria in the Metabolic Welfare of the Colonic Mucosa in Man. Gut 21, 793–798. ISSN: 0017-5749. (2021) (Sept. 1980).

31. Fetissov, S. O. Role of the Gut Microbiota in Host Appetite Control: Bacterial Growth to Animal Feeding Behaviour. Nature Reviews Endocrinology 13, 11–25. ISSN: 1759-5029, 1759-5037. (2025) (Jan. 2017).

32. Cremer, J. et al. Effect of Flow and Peristaltic Mixing on Bacterial Growth in a Gut-like Channel. Proceedings of the National Academy of Sciences of the United States of America 113, 11414–11419. ISSN: 0027-8424 (Oct. 2016).

33. Stouthamer, A. & Bettenhaussen, C. Utilization of Energy for Growth and Maintenance in Continuous and Batch Cultures of Microorganisms. Biochimica et Biophysica Acta (BBA) - Reviews on Bioenergetics 301, 53–70. ISSN: 03044173. (2025) (Feb. 1973).

34. Pirt, S. J. The Maintenance Energy of Bacteria in Growing Cultures. Proceedings of the Royal Society of London. Series B. Biological Sciences 163, 224–231. ISSN: 0080-4649, 2053-9193. (2024) (Oct. 1965).

35. Cremer, J., Arnoldini, M. & Hwa, T. Effect of Water Flow and Chemical Environment on Microbiota Growth and Composition in the Human Colon. Proceedings of the National Academy of Sciences of the United States of America 114, 6438–6443. ISSN: 0027-8424 (June 2017).

36. Popoff, M. Multifaceted Interactions of Bacterial Toxins with the Gastrointestinal Mucosa. Future Microbiology 6, 763–797. ISSN: 1746-0913, 1746-0921. (2025) (July 2011).

37. Daquigan, N., Seekatz, A. M., Greathouse, K. L., Young, V. B. & White, J. R. High-Resolution Profiling of the Gut Microbiome Reveals the Extent of Clostridium Difficile Burden. npj Biofilms and Microbiomes 3, 35. ISSN: 2055-5008. (2025) (Dec. 2017).

38. Reigadas, E. et al. Role of Clostridioides Difficile in Hospital Environment and Healthcare Workers. Anaerobe 63, 102204. ISSN: 10759964. (2025) (June 2020).

39. Dionne, L.-L. et al. Correlation between Clostridium Difficile Bacterial Load, Commercial Real-Time PCR Cycle Thresholds, and Results of Diagnostic Tests Based on Enzyme Immunoassay and Cell Culture Cytotoxicity Assay. Journal of Clinical Microbiology 51, 3624–3630. ISSN: 0095-1137, 1098-660X. (2025) (Nov. 2013).

40. Åkerlund, T., Svenungsson, B., Lagergren, Å. & Burman, L. G. Correlation of Disease Severity with Fecal Toxin Levels in Patients with Clostridium Difficile -Associated Diarrhea and Distribution of PCR Ribotypes and Toxin Yields In Vitro of Corresponding Isolates. Journal of Clinical Microbiology 44, 353–358. ISSN: 0095-1137, 1098-660X. (2025) (Feb. 2006).

41. Karlsson, S., Burman, L. G. & Åkerlund, T. Suppression of Toxin Production in Clostridium Difficile VPI 10463 by Amino Acids. Microbiology 145, 1683–1693. ISSN: 1350-0872, 1465-2080. (2025) (July 1999).

42. Merrigan, M. et al. Human Hypervirulent Clostridium Difficile Strains Exhibit Increased Sporulation as Well as Robust Toxin Production. Journal of Bacteriology 192, 4904–4911. ISSN: 0021-9193, 1098-5530. (2025) (Oct. 2010).

43. Aminzadeh, A. & Jørgensen, R. Systematic Evaluation of Parameters Important for Production of Native Toxin A and Toxin B from Clostridioides Difficile. Toxins 13, 240. ISSN: 2072-6651. (2025) (Mar. 2021).

44. Alonso, C. D. et al. Ultrasensitive and Quantitative Toxin Measurement Correlates With Baseline Severity, Severe Outcomes, and Recurrence Among Hospitalized Patients With Clostridioides Difficile Infection. Clinical Infectious Diseases 74, 2142–2149. ISSN: 1058-4838, 1537-6591. (2025) (July 2022).

45. Song, L. et al. Development and Validation of Digital Enzyme-Linked Immunosorbent Assays for Ultrasensitive Detection and Quantification of Clostridium Difficile Toxins in Stool. Journal of Clinical Microbiology 53 (ed Patel, R.) 3204–3212. ISSN: 0095-1137, 1098-660X. (2025) (Oct. 2015).

46. Sandlund, J. et al. Ultrasensitive Detection of Clostridioides Difficile Toxins A and B by Use of Automated Single-Molecule Counting Technology. Journal of Clinical Microbiology 56 (ed Onderdonk, A. B.) e00908–18. ISSN: 0095-1137, 1098-660X. (2025) (Nov. 2018).

47. Shen, Y. et al. Rapid Discrimination between Clinical Clostridioides Difficile Infection and Colonization by Quantitative Detection of TcdB Toxin Using a Real-Time Cell Analysis System. Frontiers in Microbiology 15, 1348892. ISSN: 1664-302X. (2025) (Jan. 2024).

48. Binder, H. J. Pathophysiology of Acute Diarrhea. The American Journal of Medicine 88, S2–S4. ISSN: 00029343. (2025) (June 1990).

49. Pollock, N. R. et al. Comparison of Clostridioides Difficile Stool Toxin Concentrations in Adults With Symptomatic Infection and Asymptomatic Carriage Using an Ultrasensitive Quantitative Immunoassay. Clinical Infectious Diseases 68, 78–86. ISSN: 1058-4838, 1537-6591. (2025) (Jan. 2019).

50. Wrong, O. Nitrogen Metabolism in the Gut. The American Journal of Clinical Nutrition 31, 1587–1593. ISSN: 0002-9165, 1938-3207. (2021) (Sept. 1978).

51. Liu, Y. et al. A Widely Distributed Gene Cluster Compensates for Uricase Loss in Hominids. Cell 186, 3400–3413.e20. ISSN: 00928674. (2025) (Aug. 2023).

52. Shen, T.-C. D. et al. Engineering the Gut Microbiota to Treat Hyperammonemia. Journal of Clinical Investigation 125, 2841–2850. ISSN: 0021-9738. (2022) (July 2015).

53. Rai, R., Saraswat, V. A. & Dhiman, R. K. Gut Microbiota: Its Role in Hepatic Encephalopathy. Journal of Clinical and Experimental Hepatology 5, S29–S36. ISSN: 09736883. (2025) (Mar. 2015).

54. Häberle, J. Clinical Practice: The Management of Hyperammonemia. European Journal of Pediatrics 170, 21–34. ISSN: 0340-6199, 1432-1076. (2025) (Jan. 2011).

55. Macfarlane, G. T., Cummings, J. H. & Allison, C. Protein Degradation by Human Intestinal Bacteria. Journal of general microbiology 132, 1647–1656 (June 1986).

56. You, C. et al. Coordination of Bacterial Proteome with Metabolism by Cyclic AMP Signalling. Nature 500, 301–306 (Aug. 2013).

57. Chacko, A. & Cummings, J. H. Nitrogen Losses from the Human Small Bowel: Obligatory Losses and the Effect of Physical Form of Food. Gut 29, 809–815 (June 1988).

58. Varel, V. H. & Bryant, M. P. Nutritional Features of Bacteroides Fragilis Subsp. Fragilis. Applied Microbiology 28 (1974).

59. Zeng, X. et al. Gut Bacterial Nutrient Preferences Quantified in Vivo. Cell 185, 3441–3456.e19. ISSN: 00928674. (2023) (Sept. 2022).

60. Ryvchin, R. et al. Alteration in Urease-producing Bacteria in the Gut Microbiomes of Patients with Inflammatory Bowel Diseases. Journal of Crohn’s and Colitis 15, 2066–2077. ISSN: 1873-9946, 1876-4479. (2024) (Dec. 2021).

61. Regan, M. D. et al. Nitrogen Recycling via Gut Symbionts Increases in Ground Squirrels over the Hibernation Season. Science 375, 460–463. ISSN: 0036-8075, 1095-9203. (2023) (Jan. 2022).

62. Ni, J. et al. A Role for Bacterial Urease in Gut Dysbiosis and Crohn’s Disease. Science translational medicine 9, eaah6888 (Nov. 2017).

63. Roager, H. M. et al. Colonic Transit Time Is Related to Bacterial Metabolism and Mucosal Turnover in the Gut. Nature Microbiology 1, 1–9 (June 2016).

64. Procházková, N. et al. Gut Physiology and Environment Explain Variations in Human Gut Microbiome Composition and Metabolism. Nature Microbiology 9, 3210–3225. ISSN: 2058-5276. (2025) (Nov. 2024).

65. Boekhorst, J. et al. Stool Energy Density Is Positively Correlated to Intestinal Transit Time and Related to Microbial Enterotypes. Microbiome 10, 223. ISSN: 2049-2618. (2025) (Dec. 2022).

66. Vandeputte, D. et al. Stool Consistency Is Strongly Associated with Gut Microbiota Richness and Composition, Enterotypes and Bacterial Growth Rates. Gut 65, 57–62. ISSN: 0017-5749 (Dec. 2015).

67. Falony, G. et al. Population-Level Analysis of Gut Microbiome Variation. Science 352, 560–564 (Apr. 2016).

68. Asnicar, F. et al. Blue Poo: Impact of Gut Transit Time on the Gut Microbiome Using a Novel Marker. Gut 70, 1665–1674. ISSN: 0017-5749, 1468-3288. (2025) (Sept. 2021).

69. Read, N. W., Al-Janabi, M. N., Holgate, A. M., Barber, D. C. & Edwards, C. A. Simultaneous Measurement of Gastric Emptying, Small Bowel Residence and Colonic Filling of a Solid Meal by the Use of the Gamma Camera. Gut 27, 300–308. ISSN: 0017-5749. (2025) (Mar. 1986).

70. Cummings, J. H., Jenkins, D. J. & Wiggins, H. S. Measurement of the Mean Transit Time of Dietary Residue through the Human Gut. Gut 17, 210–218 (Mar. 1976).

71. Wyman, J. B., Heaton, K. W., Manning, A. P. & Wicks, A. C. Variability of Colonic Function in Healthy Subjects. Gut 19, 146–150. ISSN: 0017-5749. (2021) (Feb. 1978).

72. Stephen, A. M. et al. The Effect of Age, Sex and Level of Intake of Dietary Fibre from Wheat on Large-Bowel Function in Thirty Healthy Subjects. British Journal of Nutrition 56, 349–361. ISSN: 0007-1145, 1475-2662. (2022) (Sept. 1986).

73. Stephen, A. M. & Cummings, J. H. The Microbial Contribution to Human Fecal Mass. Journal of Medical Microbiology 13, 45–56. ISSN: 0022-2615, 1473-5644. (2021) (Feb. 1980).

74. Abrahamsson, H. & Antov, S. Accuracy in Assessment of Colonic Transit Time with Particles: How Many Markers Should Be Used?: Colonic Transit Time. Neurogastroenterology & Motility 22, 1164–1169. ISSN: 13501925. (2025) (Nov. 2010).

75. Abrahamsson, H., Antov, S. & Bosaeus, I. Gastrointestinal and Colonic Segmental Transit Time Evaluated by a Single Abdominal X-ray in Healthy Subjects and Constipated Patients. Scandinavian Journal of Gastroenterology 23, 72–80. ISSN: 0036-5521, 1502-7708. (2025) (Jan. 1988).

## References

1. Penry, D. L. & Jumars, P. A. Modeling Animal Guts as Chemical Reactors. The American Naturalist 129, 69–96. ISSN: 0003-0147, 1537-5323. (2025) (Jan. 1987).

2. Martìnez Del Rio, C., Cork, S. J. & Karasov, W. H. in The Digestive System in Mammals (eds Chivers, D. J. & Langer, P.) 1st ed., 25–53 (Cambridge University Press, July 1994). ISBN: 978-0-521-44016-5 978-0-521-02085-5 978-0-511-66171-6. (2025).

3. Caton, J. M. & Hume, I. D. Chemical Reactors of the Mammalian Gastro-Intestinal Tract. Zeitschrift für Säugetierkunde 65, 33–50. ISSN: 0044-3468 (2000).

4. Le Feunteun, S. et al. Physiologically Based Modeling of Food Digestion and Intestinal Microbiota: State of the Art and Future Challenges. An INFOGEST Review. Annual Review of Food Science and Technology 12, 149–167. ISSN: 1941-1413, 1941-1421. (2025) (Mar. 2021).

5. Labarthe, S. et al. A Mathematical Model to Investigate the Key Drivers of the Biogeography of the Colon Microbiota. Journal of Theoretical Biology 462, 552–581. ISSN: 00225193. (2025) (Feb. 2019).

6. Muñoz-Tamayo, R., Laroche, B., Walter, É., Doré, J. & Leclerc, M. Mathematical Modelling of Carbohydrate Degradation by Human Colonic Microbiota. Journal of Theoretical Biology 266, 189–201 (Sept. 2010).

7. Taghipoor, M., Lescoat, P., Licois, J.-R., Georgelin, C. & Barles, G. Mathematical Modeling of Transport and Degradation of Feedstuffs in the Small Intestine. Journal of Theoretical Biology 294, 114–121. ISSN: 00225193. (2025) (Feb. 2012).

8. Codutti, A., Cremer, J. & Alim, K. Changing Flows Balance Nutrient Absorption and Bacterial Growth along the Gut. Physical Review Letters 129, 138101. ISSN: 0031-9007, 1079-7114. (2025) (Sept. 2022).

9. Salari, A. & Cremer, J. Diurnal Variations in Digestion and Flow Drive Microbial Dynamics in the Gut. PRX Life 3, 023012. ISSN: 2835-8279. (2025) (June 2025).

10. Arnoldini, M., Cremer, J. & Hwa, T. Bacterial Growth, Flow, and Mixing Shape Human Gut Microbiota Density and Composition. Gut Microbes, 1–8 (July 2018).

11. Arnoldini, M. et al. Quantifying the Varying Harvest of Fermentation Products from the Human Gut Microbiota. Cell 188, pP5332–5342.E16. ISSN: 0092-8674. (2025) (Sept. 2025).

12. Fromentin, S. et al. Microbiome and Metabolome Features of the Cardiometabolic Disease Spectrum. Nature Medicine 28, 303–314. ISSN: 1078-8956, 1546-170X. (2025) (Feb. 2022).

13. Vieira-Silva, S. et al. Statin Therapy Is Associated with Lower Prevalence of Gut Microbiota Dysbiosis. Nature 581, 310–315. ISSN: 0028-0836, 1476-4687. (2025) (May 2020).

14. Forslund, S. K. et al. Combinatorial, Additive and Dose-Dependent Drug–Microbiome Associations. Nature 600, 500–505. ISSN: 0028-0836, 1476-4687. (2025) (Dec. 2021).

